## Supplementary Material for "*Cis*-regulatory modes of *Ultrabithorax* inactivation in butterfly forewings"

### SUPPLEMENTARY INFORMATION

| Species | sgRNA name | Target Sequence (5' to 3')<br>PAM sequence not shown |
|---|---|---|
| *J. coenia* | *Antp-Ubx_BE* | CTCGAATATGGAGATATCGG |
| | *UbxCRE11a3* | ACGGACCTCCGCTTTCCTGG |
| | *UbxCRE11c6* | AACTGGTGCAGTGCCTTGTA |
| *J. coenia* + *V. cardui* | *UbxCRE11a2* | CTACTCTGTTCGGACATTCG |
| | *UbxCRE11c5* | GCTGCCGCGAGTCTGAATCG |
| | *UbxCRE11b9* | TTCATGTATGAACCATGACG |
| | *UbxIT1#1* | CCTTCGCATAAGTTCGGATAGG |
| | *Bxd1* | TATCGGTCGTTCGTCACACA |
| *V. cardui* | *UbxIT1#2* | CTCGGCTATGTGTCGAGGGC |

**Table S1. List of sgRNAs used in CRISPR experiments.**

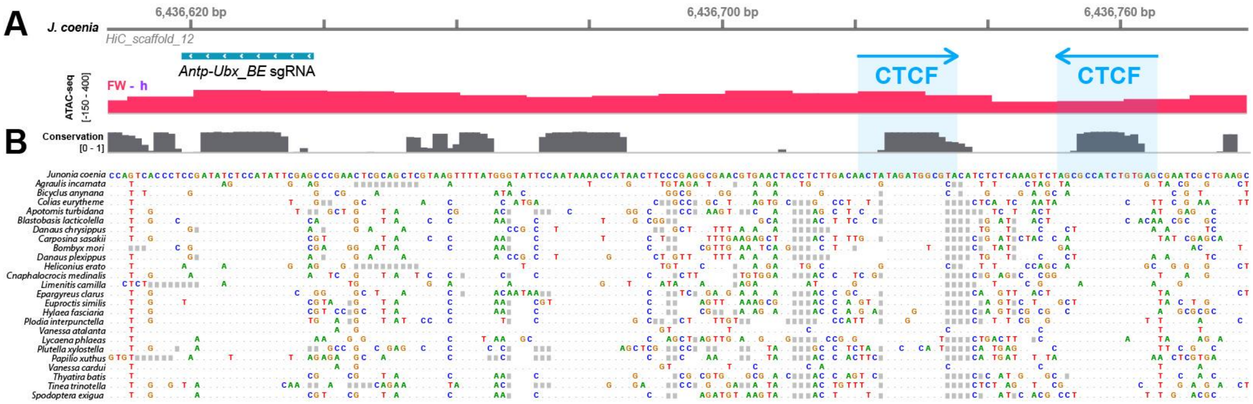


**Figure S1. Prediction of two conserved CTCF binding sites at *Antp-Ubx_BE*.** (**A**) Sequence-level view of a 180-bp genomic interval including the *Antp-Ubx_BE* sgRNA (turquoise) in *J. coenia*, overlapping with an ATAC-seq peak of forewing-enriched chromatin opening (red). The CRISPR target is about 100 bp away from two predicted binding sites for the *Drosophila* CTCF insulator protein. (**B**) High-level of nucleotide conservation at thesgRNA site and CTCF motifs across Lepidoptera and Trichoptera representative genomes, indicative of functional constraints on these sequences.

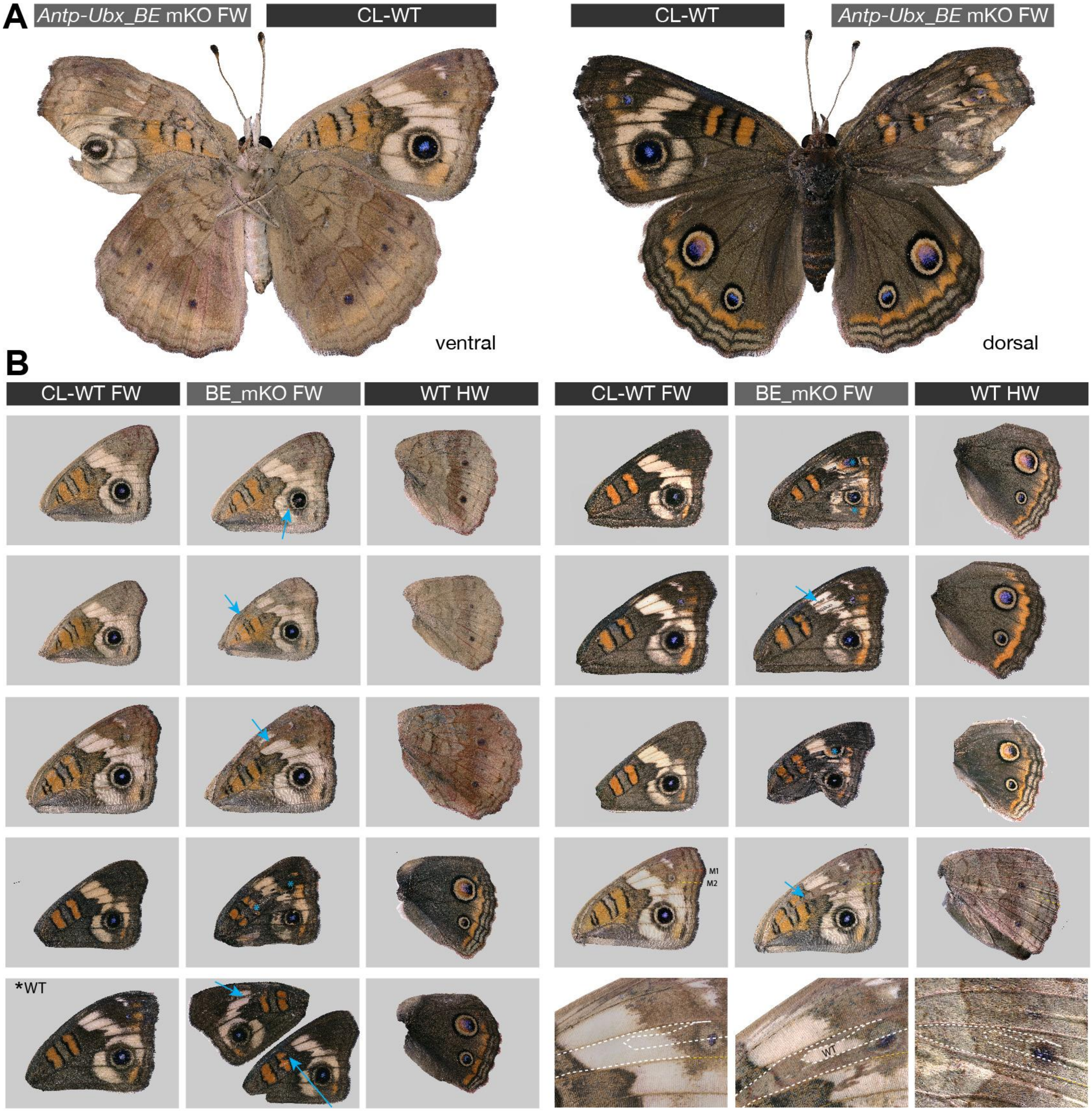


**Figure S2. CRISPR perturbation of the *Antp-Ubx* boundary element results in FW-to-HW homeosis.** (**A**) Example of an *Antp-Ubx_BE* crispant with a unilateral phenotype on the right forewing. (**B**) Additional examples of forewing homeoses in *Antp-Ubx_BE* crispant. Wing sets (forewing mKO mutants and corresponding contralateral WT) are shown with one of the wings horizontally flipped to show the mutant wings in left-to-right orientation. Arrows : small mutant clones. Asterisks : large mutant clones.

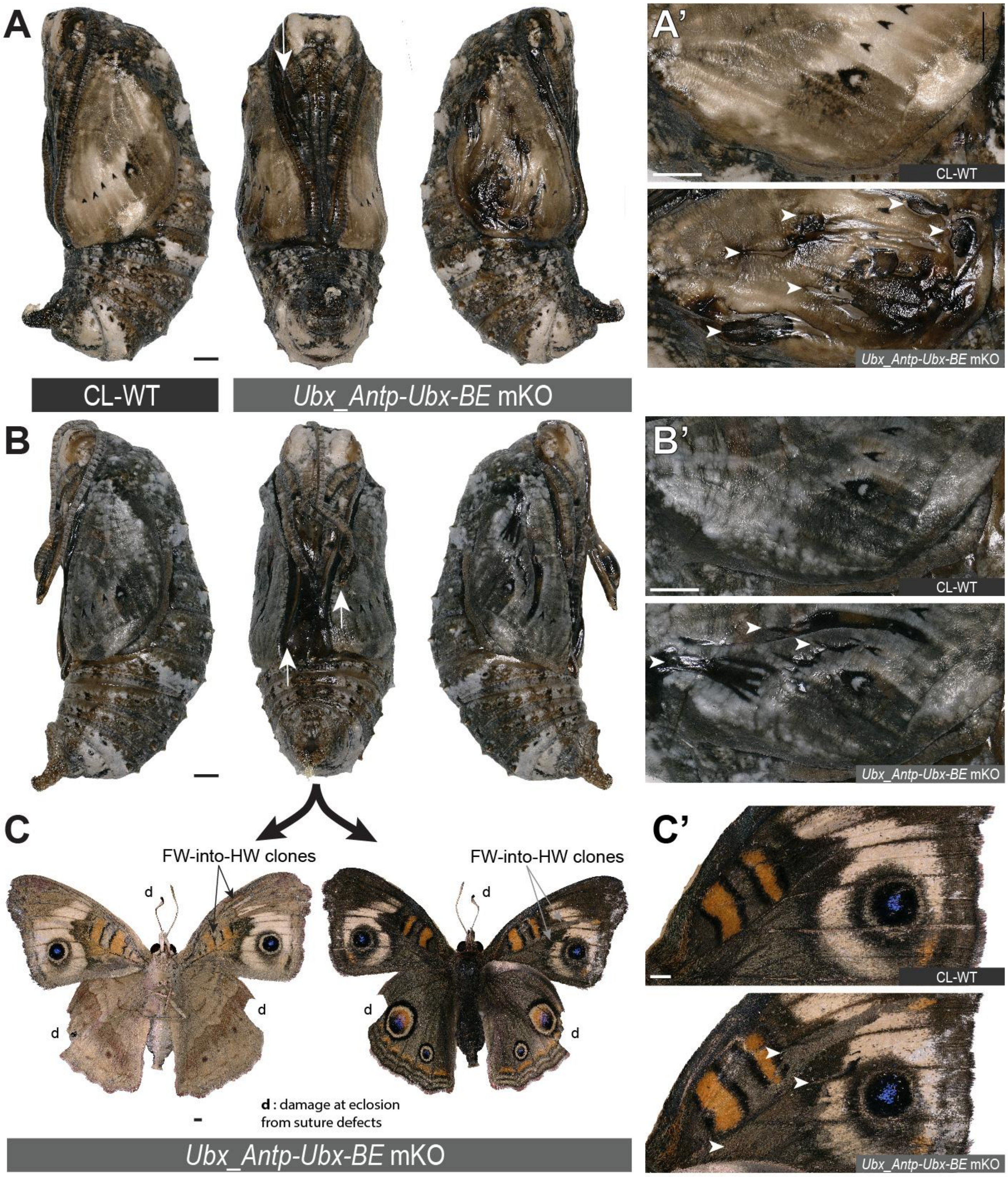


**Figure S3. Pupal defects following FW→HW homeosis in *Antp-Ubx_BE* crispants.** (**A-B**) Contralateral (CL) and, forewing mosaic knockout (mKO) mutants following CRISPR targeting of *Antp-Ubx_BE* in *J. coenia*. The two pupae show suture defects in the midline appendages (arrows). (**A'-B'**) Magnified views of the crispant forewings, showing defective cuticle (arrowheads). (**C-C'**) Crispant adult butterfly emerged from the pupa in panel B. White arrowheads in C’ highlight the match between dorsal forewing clones and the pupal forewing cuticle defects shown in B’. Scale bars : 1 mm.

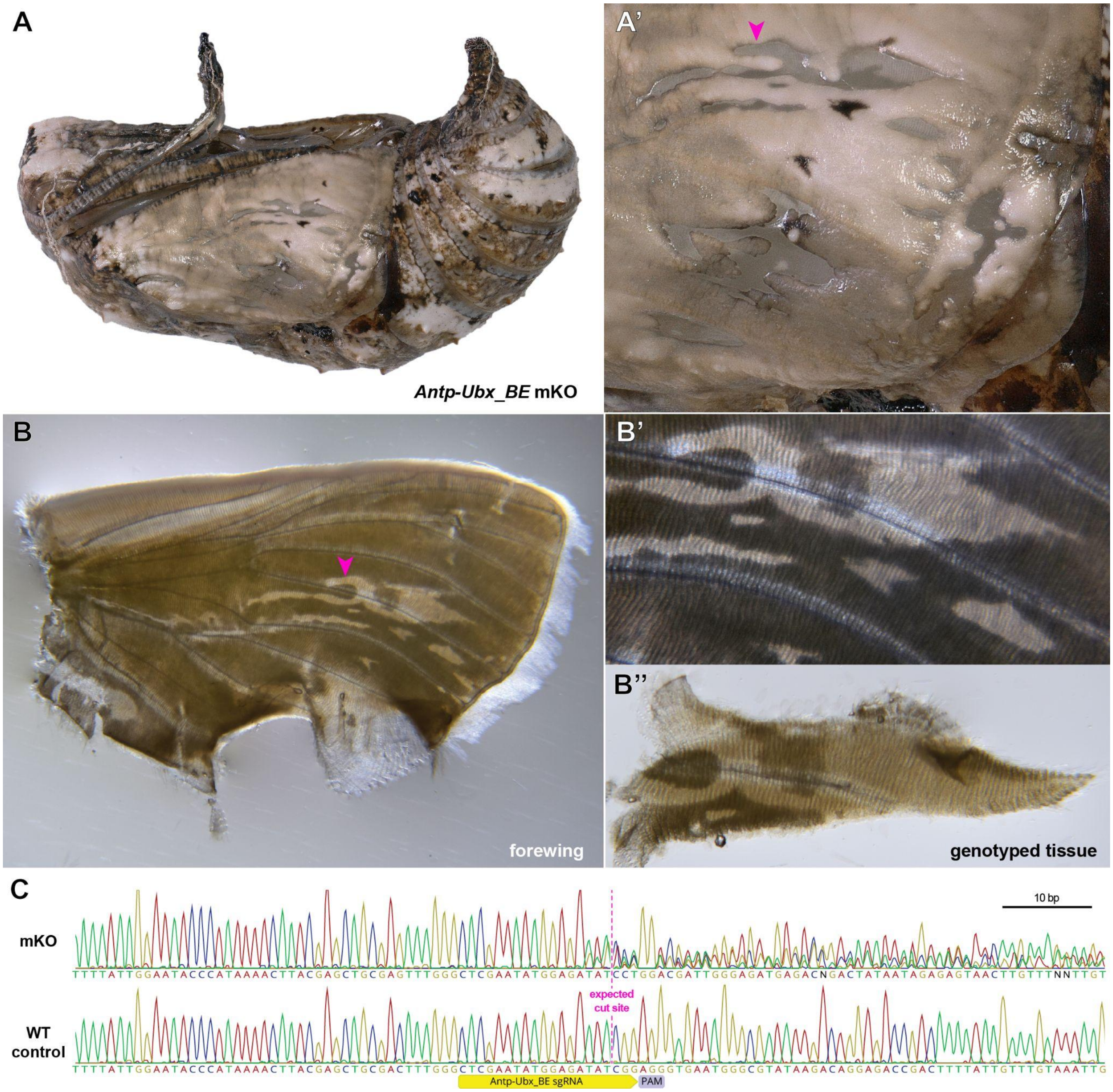


**Figure S4. Validation of CRISPR-induced DNA lesions in an *Antp-Ubx_BE* crispant pupal forewing.** (**A-A’**) Pupal forewing cuticle phenotype of an *Antp-Ubx_BE J. coenia* crispant, as in Fig. S3. (**B-B’’**) Aspect of the same forewing under trans-illumination following dissection out of the pupal case. Regions from mutant clones have a more transparent appearance. (**C**). Sanger sequencing of an amplicon targeting the *Antp-Ubx_BE* region in the mutant tissue shown in panel B”, compared to a control wing tissue, showing mixed chromatogram around the expected CRISPR cutting site due to indel mutations from non-homologous end-joining.

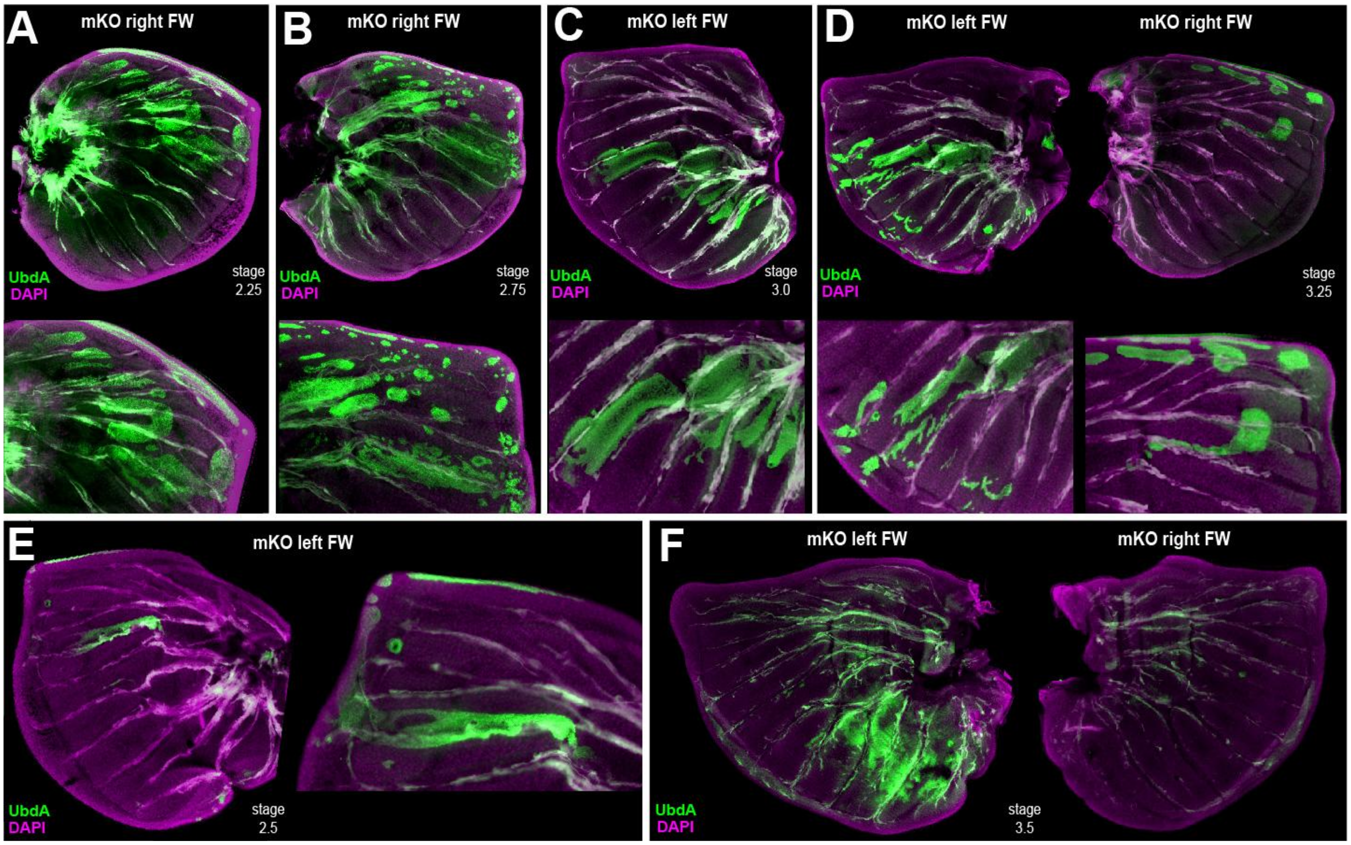


**Figure S5. Additional examples of ectopic UbdA and FW→HW homeosis in *Antp-Ubx_BE* crispant larval forewings.** (**A-F**) Each panel shows forewings with ectopic detection of UbdA (FP6.87 monoclonal antibody, green), dissected at the fifth instar stage. Panels D and F are wing sets from individual crispants. Panels E and C are mutant contralateral wings of the mutant forewings shown in Figs. 4D and E, respectively.

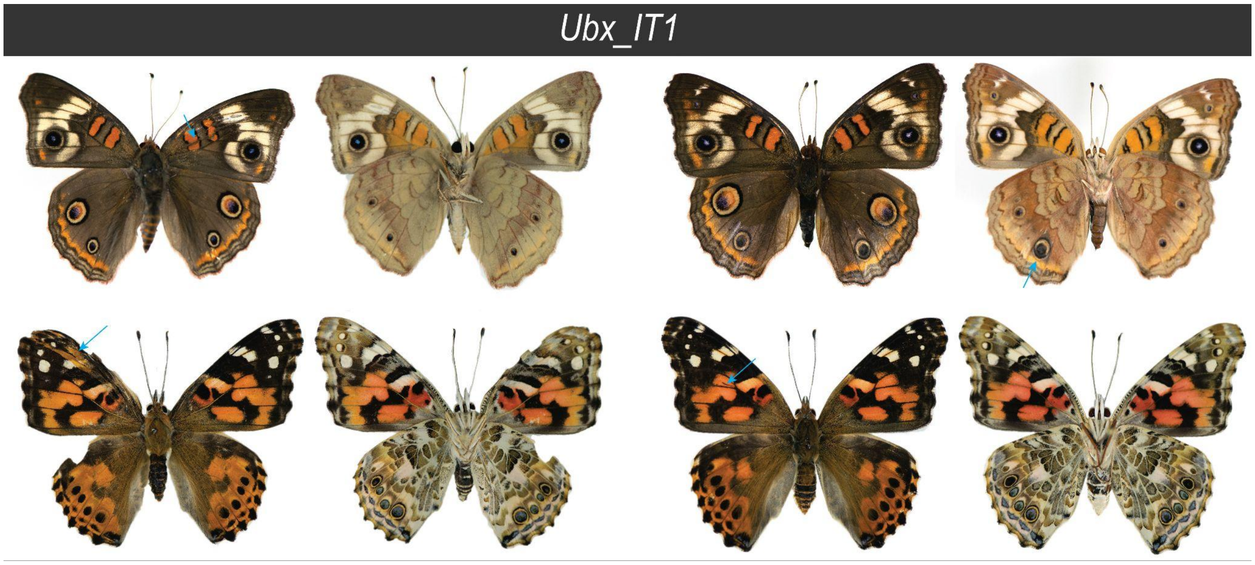


**Figure S6. Additional mutant phenotypes from CRISPR-mediated interrogation of *lncRNA_Ubx-IT1* 5' region in *J. coenia* (top) and *V. cardui* (bottom).** Cyan arrows : mutant clones.

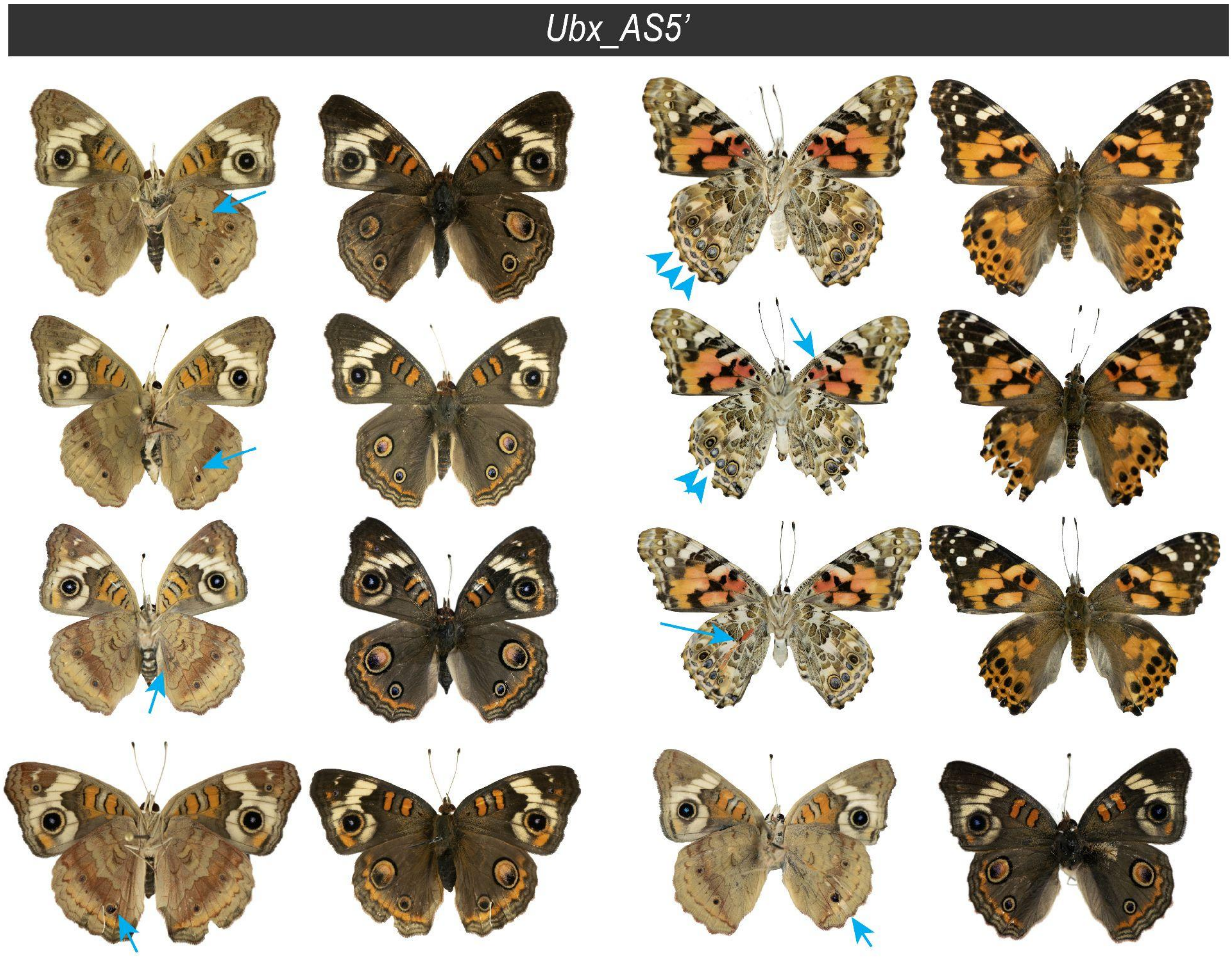


**Figure S7. Additional mutant phenotypes from CRISPR-mediated interrogation of the *lncRNA_Ubx-AS*5' region in *J. coenia* and *V. cardui*.** Arrows : mutant clones. Arrowheads : white eyespot foci.

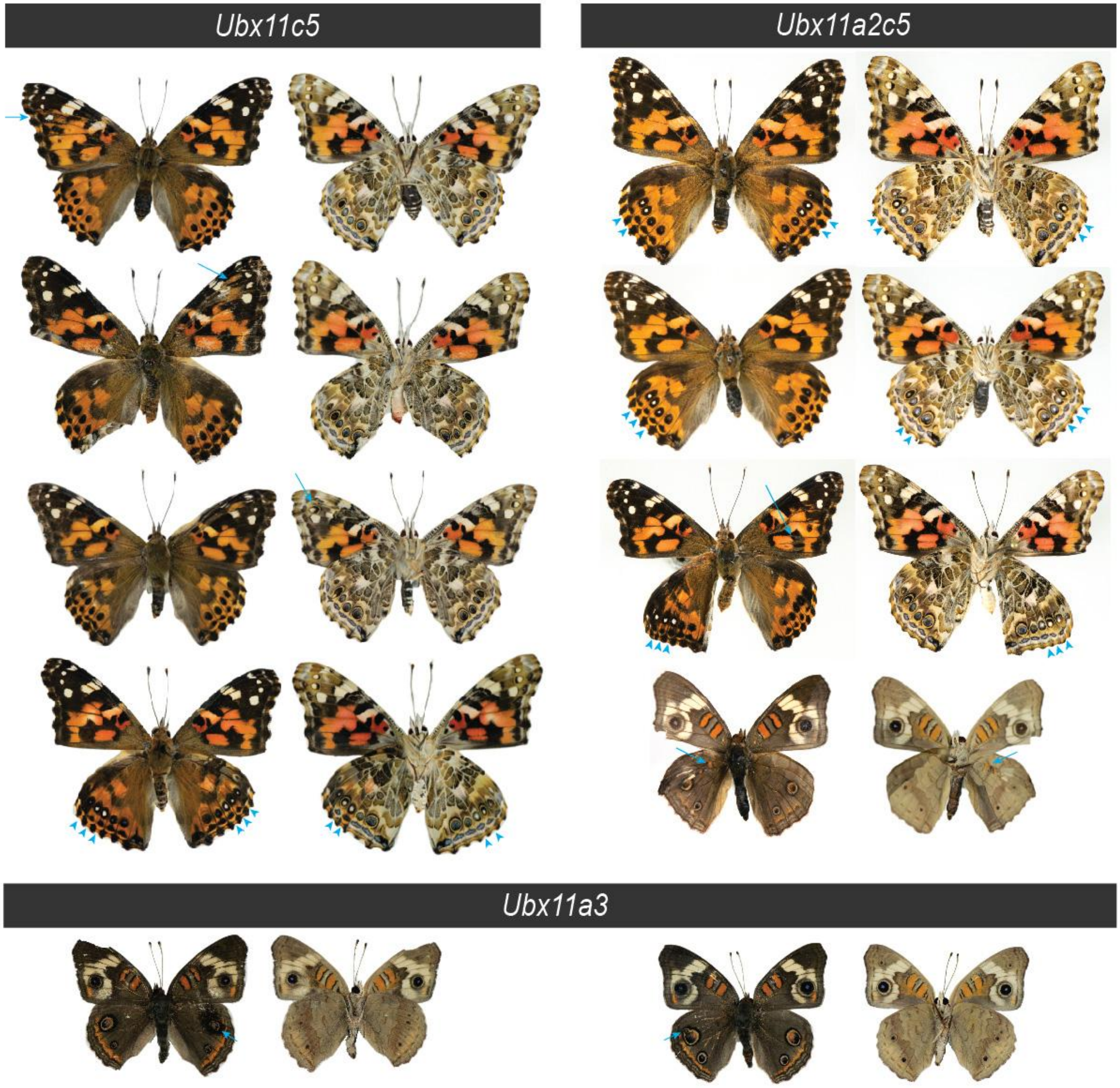


**Figure S8. Additional mutant phenotypes from CRISPR-mediated interrogation of *CRM11* in *J. coenia* and *V. cardui* show bidirectional homeoses and non-homeotic eyespot changes.** Arrows : mutant clones. Arrowheads : white eyespot foci.

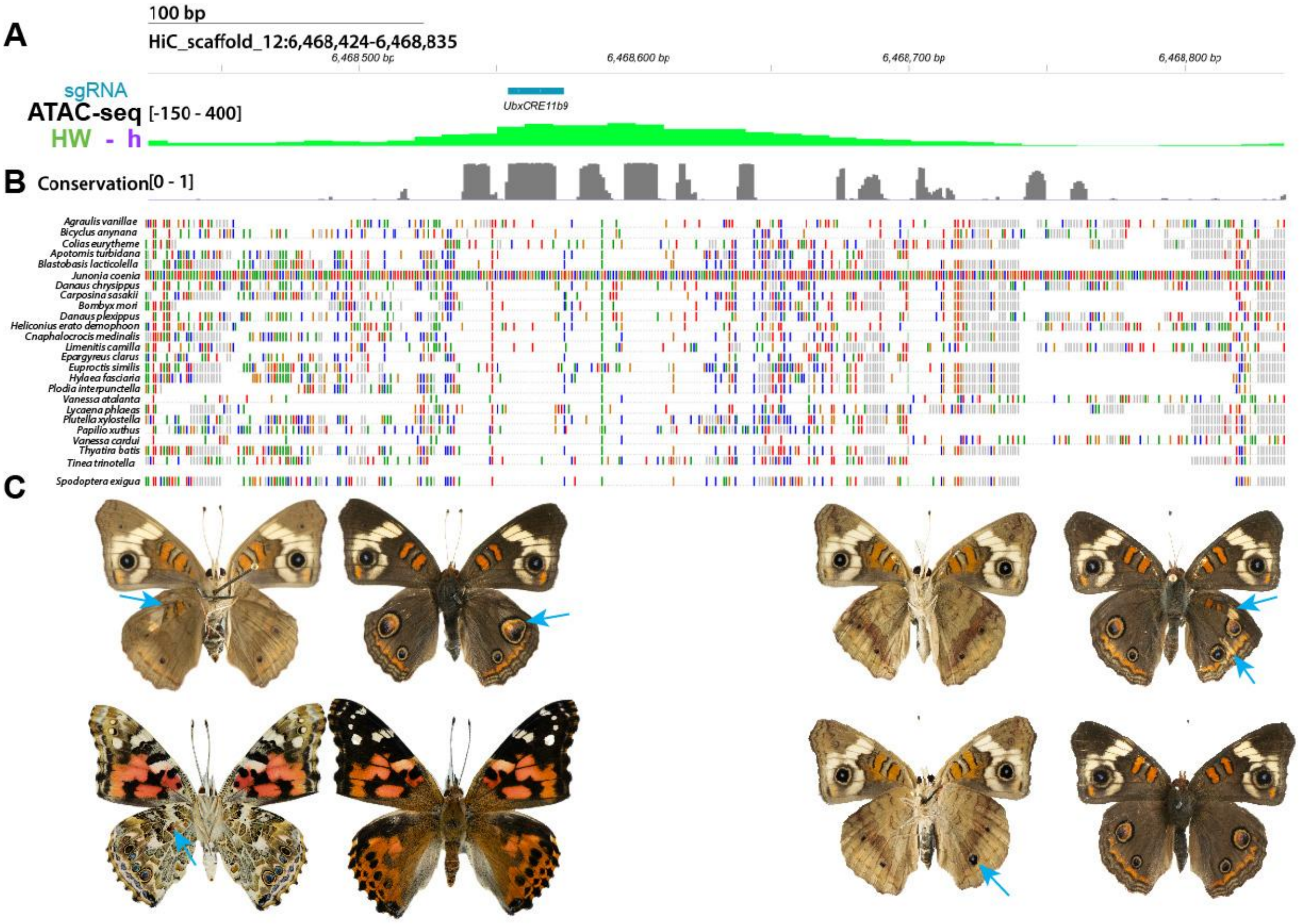


**Figure S9. CRISPR perturbation of the conserved *Ubx_CRE11b* results in HW→FW homeoses**. (**A-B**) The *UbxCRE11b9* sgRNA targets a hindwing-enriched ATAC peak with strong conservation across genomes from 23 Lepidoptera and 2 Trichoptera species (gray : PhastCons scores). Colored bars denote variation from the *J. coenia* reference. (**C**) *Jc_UbxCRE_11b9* crispant butterflies exclusively showed HW→FW transformed clones (blue arrows in both *J. coenia* and *V. cardui*).

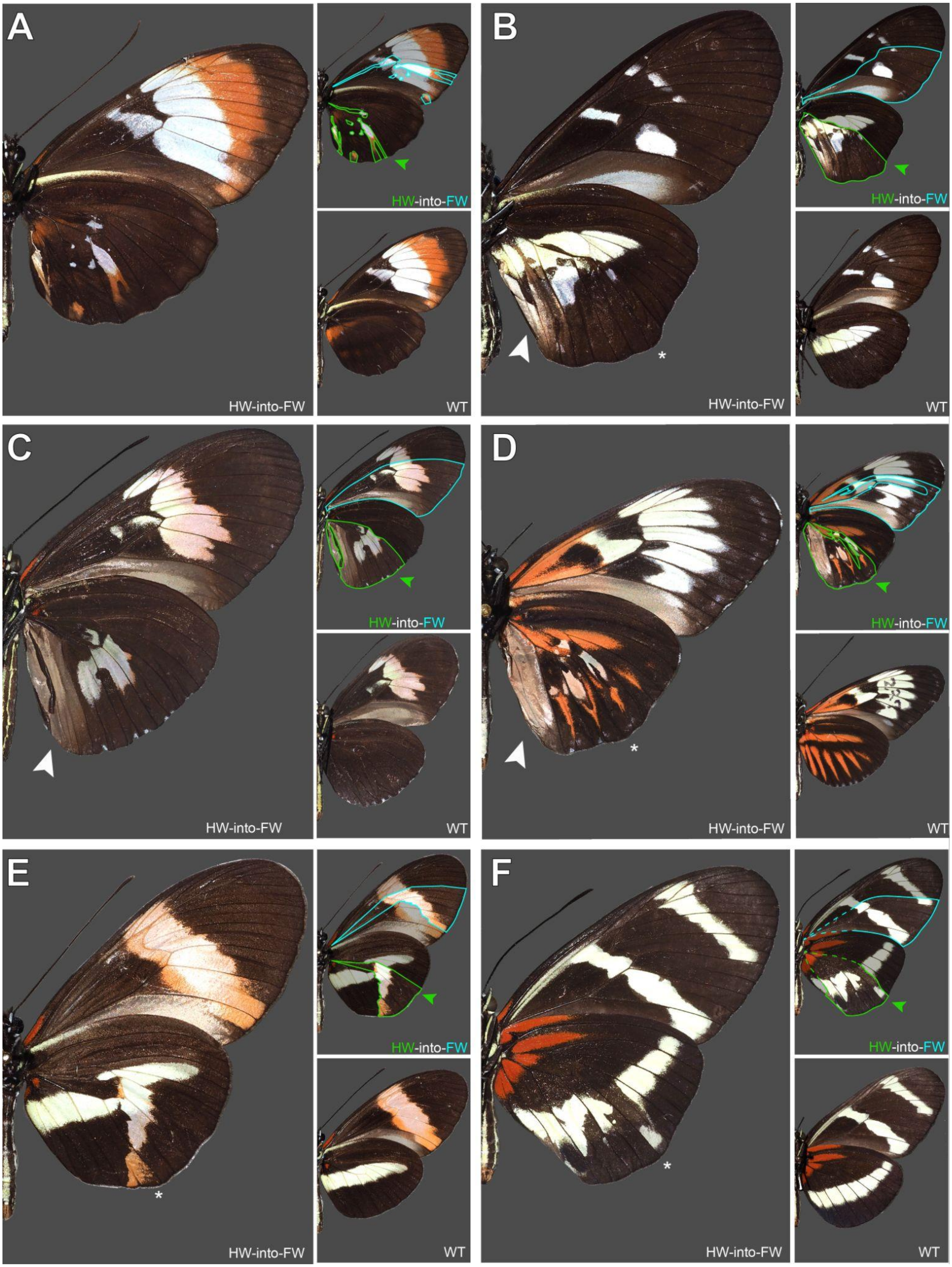
A
HW-into-FW
WT
B
HW-into-FW
WT
C
HW-into-FW
WT
D
HW-into-FW
WT
E
HW-into-FW
WT
F
HW-into-FW
WT

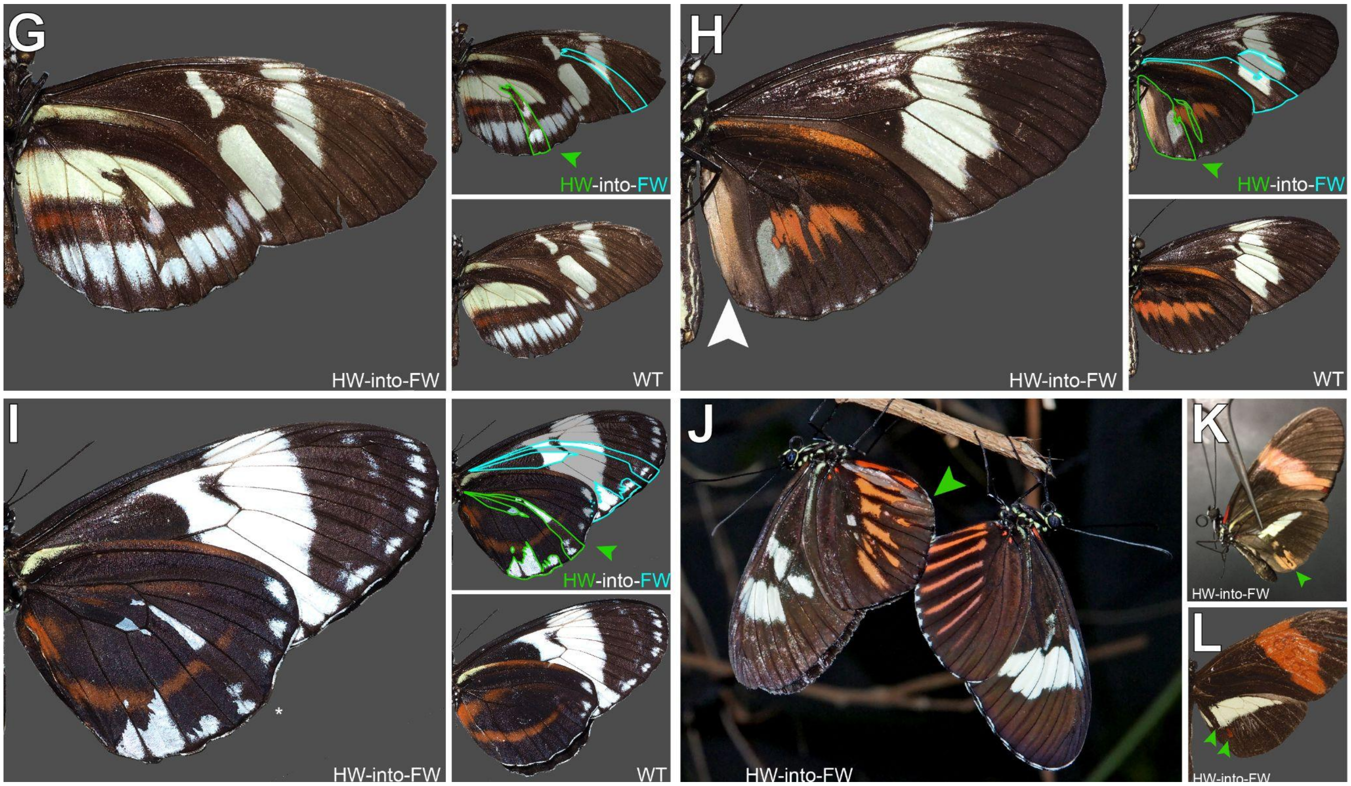


**Figure S10 (previous page, continued above).** Hindwing homeoses in *Heliconius* butterfly spontaneous mutants from pure stocks, hybrid cultures and wild-caught individuals from the L.E. Gilbert collection (UT Austin). White arrowheads: homeotic clones including the acquisition of ventral forewing coupling scales. Asterisks : local deformation of hindwings relative to wild-type. All hindwing homeoses are ventral except in panel L. **A.** *Heliconius cydno galanthus x H. melpomene rosina* (Costa Rica), cross J31, August 1987. **B.** *Heliconius cydno gustavi*, captive stock from Saladito (Colombia), September 1991. **C.** *Heliconius melpomene madeira* (Brazil) *x Heliconius melpomene plesseni* (Ecuador), September 2012. **D.** *H. m. rosina* (Costa Rica) *x Heliconius melpomene madeira* (Brazil) *x H. cydno galanthus* (Costa Rica) mixed population, December 2015. **E.** *H. m. rosina,* captive stock from Osa Peninsula (Costa Rica), September 1991. **F.** *Heliconius hewitsoni,* captive stock from Osa Peninsula (Costa Rica), July 2005. **G.** *Heliconius cydno cydnides*, captive stock from natural hybrid zone in Dagua Pass (Colombia), May 1989. **H.** *H. m. rosina* (Costa Rica) *x H. m. madeira* (Brazil) *x H. c. galanthus* (Costa Rica) mixed population, June 2016. **I.** *H. c. galanthus x H. m. rosina* crossed three times, and back to *H. c. galanthus*, August 2014. **J.** *Heliconius melpomene malleti* (Ecuador) x *H. m. plesseni* (Ecuador) hybrid stock, 2010. **K.** *H. m. rosina* captive stock, Costa Rica. **L.** *H. m. rosina* captive stock, Osa Peninsula (Costa Rica), March 1987, in dorsal view.
